# Viral kinetics of H5N1 infections in dairy cattle

**DOI:** 10.1101/2025.02.01.636082

**Authors:** Oliver Eales, James M. McCaw, Freya M. Shearer

**Affiliations:** Infectious Disease Dynamics Unit, Centre for Epidemiology and Biostatistics, Melbourne School of Population and Global Health, The University of Melbourne, Australia; School of Mathematics and Statistics, The University of Melbourne, Australia; Infectious Disease Ecology and Modelling, The Kids Research Institute, Perth, Australia

## Abstract

Since early-2024 unprecedented outbreaks of highly pathogenic avian influenza H5N1 clade 2.3.4.4b have been ongoing in dairy cattle in the United States with significant consequences for the dairy industry and public health. Estimation of key epidemiological parameters is required to support outbreak response, including predicting the likely effectiveness of interventions and testing strategies. Here we pool limited publicly available data from three studies of naturally and experimentally infected dairy cattle. We quantify Ct value trajectories of infected dairy cattle and the relationship between Ct value and the log-titre of infectious virus, β_2_ a proxy for infectiousness. We estimate that following infection peak Ct values are rapidly reached within 1–2 days with a population mean Ct value of 16.9 (13.2, 20.5). We identify a critical threshold Ct value of 21.5 (20.1, 23.6), with values of Ct value above this threshold representing little-to-no infectious viral load. Finally, we estimate the distribution of the duration of infectiousness for dairy cattle (i.e. the duration their Ct value remains above the critical threshold) with a population median of 6.2 (2.8, 13.1) days.

## Introduction

Since early-2024 unprecedented outbreaks of highly pathogenic avian influenza H5N1 clade 2.3.4.4b (genotype B3.13) have been ongoing in dairy cattle in the United States, following its introduction from wild birds [1]. As of 2 February 2025, 951 cases have been confirmed across 16 states [2]. These outbreaks have significant consequences for the dairy industry and public health. Reduced milk output of infected cattle, among other clinical symptoms [3], is resulting in economic loss to the dairy industry. Spillover infections in dairy farm workers and raw milk contamination highlight the increasing public health risks of H5N1 clade 2.3.4.4b (genotype B3.13). Additionally, there is concern that continued circulation of H5N1 strains between cattle and other mammals may result in the virus adapting for human-to-human transmission [4], increasing the risk of a future influenza pandemic. To minimise these risks it is critical that outbreaks are: detected early through effective testing strategies; and rapidly contained through the implementation of effective outbreak controls.

Information on key epidemiological parameters is required to inform effective testing and outbreak control strategies. For example, the duration of infectiousness, the timing of the onset of infectiousness relative to clinical symptoms, and the timing of infection detection and isolation, all strongly influence the effectiveness of case isolation in controlling an outbreak [5]. Additionally, the effectiveness of testing of pooled samples — a method for testing entire dairy herds with limited resources — will strongly depend on the sensitivity of a test for detecting viral RNA, within-host viral kinetics, and the type of sample collected (e.g. blood, serum, urine). Previous studies have suggested that milk and milking procedures are the dominant route of cow-to-cow transmission, with the highest viral loads detected in milk samples from infected cattle [3]. Thus quantifying the viral kinetics of H5N1 2.3.4.4b (genotype B3.13) in milk samples of infected dairy cattle can inform testing and other outbreak control strategies.

Here we have pooled the limited publicly available data from three recently published studies [3,6,7] and performed statistical analyses to quantify H5N1 viral kinetics in milk samples of infected dairy cattle. We estimate the trajectories of cycle threshold (Ct) values as measured through qRT-PCR testing of milk samples (a proxy for viral load), the relationship between Ct value and the log-titre of infectious virus (a proxy for infectiousness), and the duration of infectiousness for dairy cattle (i.e. the duration that their Ct value remains above a critical threshold). We interpret our findings, with respect to their implications for testing strategies and controllability of outbreaks in dairy cattle.

### Quantifying Ct value trajectories

We developed a Bayesian hierarchical model for quantifying Ct value trajectories in the milk of infected dairy cattle (see Methods for full description), building on models used to infer Ct value trajectories of SARS-CoV-2 infections during the COVID-19 pandemic [8,9]. Ct values are modelled as a function of time since infection, with parameters describing the peak viral load (i.e. the minimum Ct value reached, as low Ct values correspond to high viral loads), the time since infection at which the peak viral load is reached, and two rate parameters describing the initial viral growth rate before the peak (decline in Ct value), and the viral decay rate following the peak (increase in Ct value). We estimate population mean parameters and variation in these parameters between individuals (standard deviation in the mean parameters). The model is fit to longitudinal data from five experimentally infected dairy cattle (Fig.1a) with daily RT-PCR testing of milk samples [6,7], and 12 naturally infected dairy cattle (Fig.1b) who were tested on days 3, 16, and 31 post clinical diagnosis [3]. We assumed that the naturally infected dairy cattle were already in a phase of viral decay.

**Figure 1:**
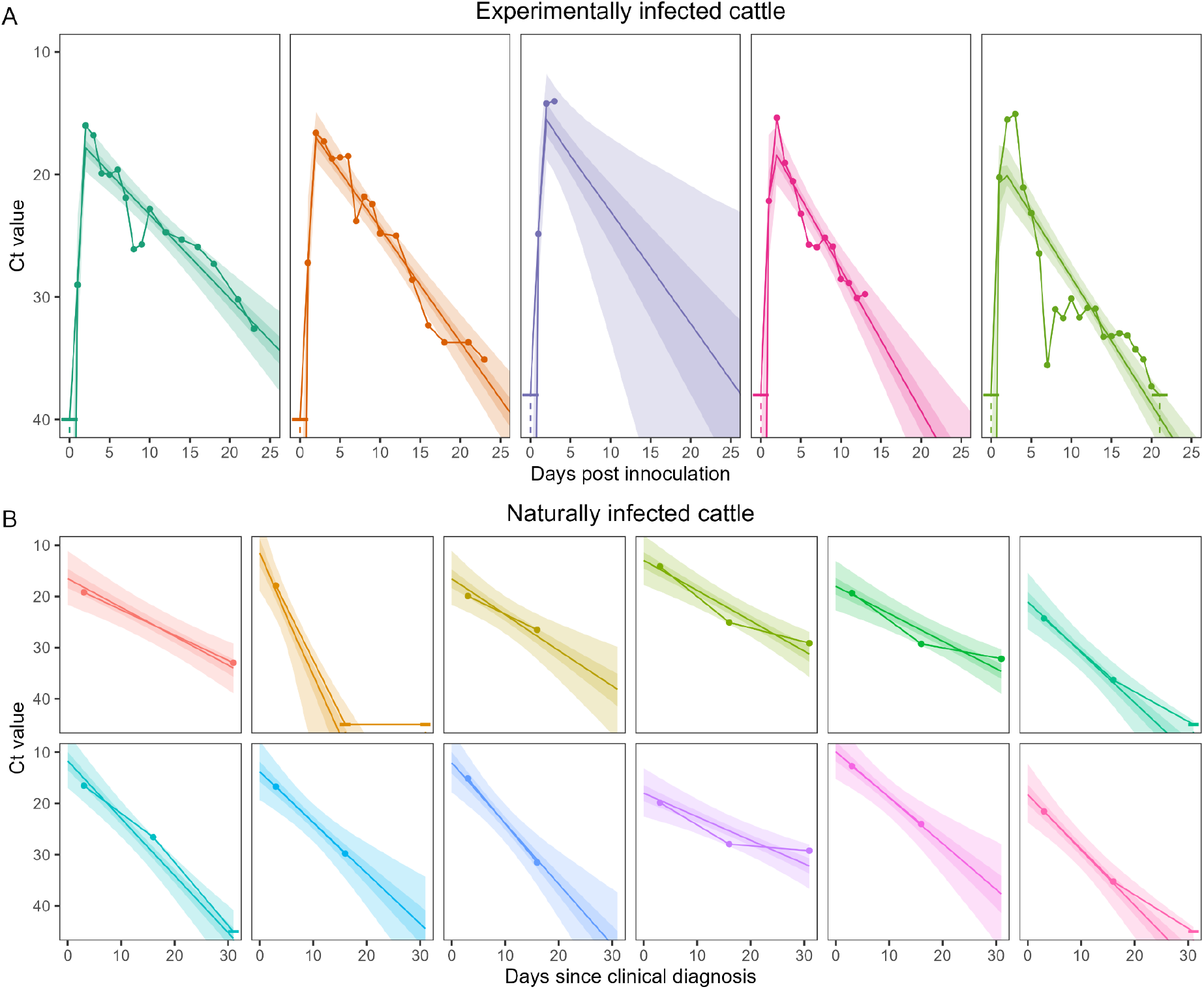
Ct value trajectory model posterior fit to data. Ct value trajectory model posterior estimates of the individual-level Ct value trajectories for experimentally infected cattle (A) and naturally infected cattle (B). Ct value trajectory posterior estimates are shown with median (line), and 50% (dark shaded region) and 95% (light shaded region) credible intervals. Data points are connected via solid lines, with uncensored data shown as solid points, and censored data shown as a horizontal dash with a dashed line extending to the possible range of values. The data of the first two (dark green, orange) experimentally infected cows were from Baker et al. 2024 [7], and the data of the final free experimentally infected cows were from Halwe et al. 2024 [6]. The data for naturally infected cows were from Caserta et al. 2024 [3].

We estimate that following infection minimum Ct values in milk are rapidly reached within 1–2 days with a population mean of 1.10 (0.84, 1.34) days (Fig.2a, SFig.1, SFig.2). We estimate a population mean in minimum Ct value of 16.9 (13.2, 20.5), followed by a gradual decline in viral load, with a population mean increase in Ct value of 0.88 (0.67, 1.18) per day. Estimates of the variation in parameters between individuals were uncertain, reflecting the small sample size of animals for which an entire Ct trajectory was available. However, there was greater certainty in our estimates for the variation in the viral decay rate, due to the additional data describing natural infections. Estimated variability in the viral decay rate suggests heterogeneity in the duration that an animal’s viral load remains elevated (Fig 2A). We performed a sensitivity analysis fitting the model to subsets of the data (see Methods) and found no evidence of different parameter values between experimentally and naturally infected cattle (SFig.1), although there were (as expected) higher levels of uncertainty in estimates.

**Figure 2:**
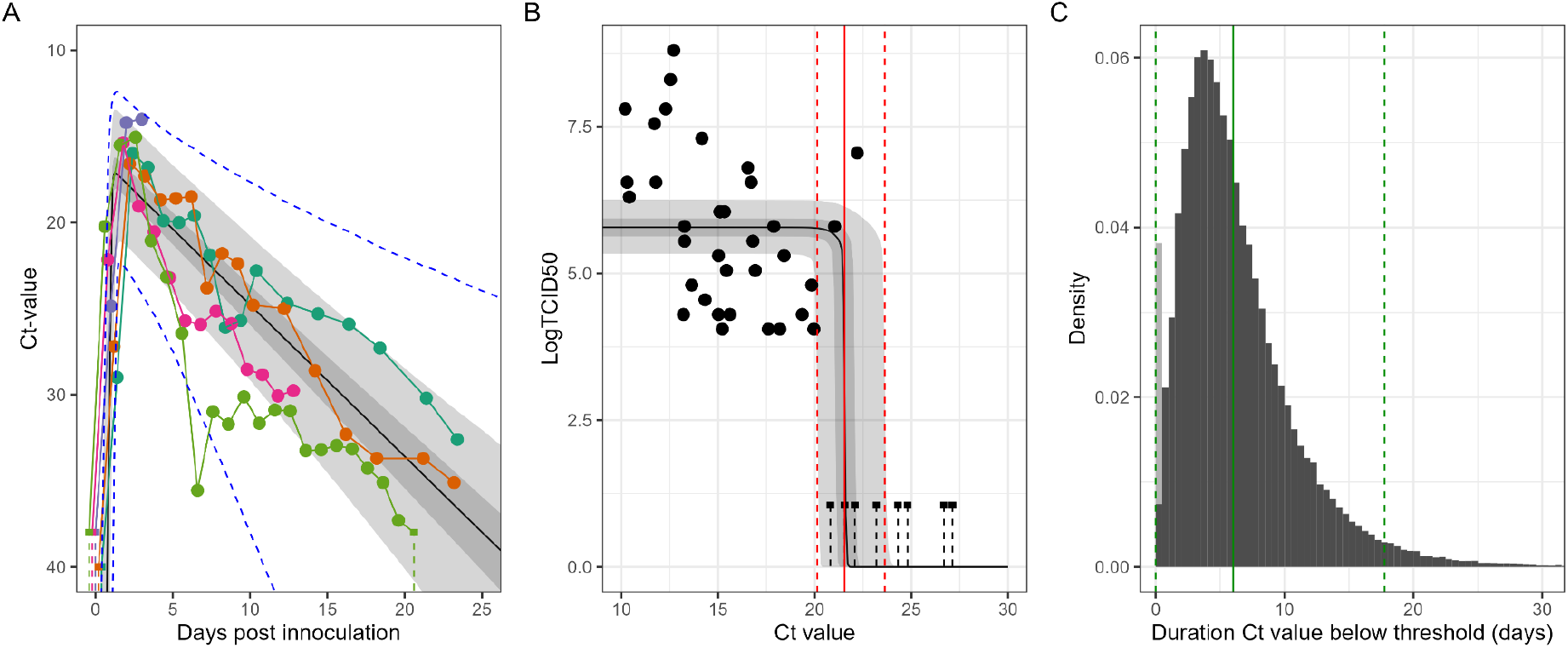
Population distribution of Ct value trajectories and infectiousness. **(A)** Ct value trajectory model posterior estimates of the population mean Ct value trajectory (i.e. using population mean parameters). Posterior estimates are shown with median (line), and 50% (dark shaded region) and 95% (light shaded region) credible intervals. Longitudinal data (points connected by solid lines) for the experimentally infected cows [6,7] are also shown (colours match Fig.1) with censored data represented by solid squares with a dashed line extending to the possible range of values. The central 95% interval (dashed blue lines) of the estimated population distribution of Ct value trajectories using the median of the population parameters (means and standard deviations) is also shown. **(B)** Posterior estimates of the relationship between log titre of infectious virus and Ct value are shown with median (line), and 50% (dark shaded region) and 95% (light shaded region) credible intervals. Uncensored data are shown as solid points, and censored data it shows as solid squares with a dashed line extending to the possible range of values. Also shown is the median (solid red line) and 95% credible interval (dashed red lines) in the threshold Ct value at which the log-titre of infectious virus rapidly increases from 0 to a maximum (parameter in methods). **(C)** The distribution of the duration that individual Ct value trajectories remain below the critical threshold Ct value (when they are likely infectious) estimated using: median population parameters (means and standard deviations) from the Ct value trajectory model (panel A); and the median threshold Ct value (solid red line in panel B). Also shown is the mean of the distribution (solid green line) and central 95% interval of the distribution (dashed green lines). The portion of the distribution that has a value of 0 is shaded light grey.

### Pooled testing of milk vats

Timely detection of H5N1 outbreaks in dairy farms is crucial for minimising transmission within a farm and the risk of spread between farms. Early detection can allow rapid implementation of control measures such as case isolation, quarantine, livestock transport controls, and enhanced cleaning protocols. While it would likely be impractical to frequently test all cattle — or even all clinically affected cattle — pooled testing of samples from multiple cattle could be feasible and effective at identifying outbreaks. However, the utility of pooled testing depends on the effectiveness of the test at identifying an infection in a diluted sample (i.e. diluted by many uninfected samples). Pooled testing of milk samples is an obvious strategy to employ for detection of H5N1 outbreaks in dairy farms: the highest Ct values have consistently been measured in milk samples of infected animals; daily (at least) milking is routine; and milk samples for a single dairy farm are often pooled into a central milk vat.

We calculated the expected Ct value of a pooled milk sample containing the milk from a single infected animal, varying the: Ct value of the milk of the infected animal if tested in isolation; the number of milk samples in the pooled sample; and the efficiency of the rt-PCR test used to measure the Ct value (Fig 3). The ability of the pooled test to identify the presence of a single infected animal will depend on the relative value of the expected Ct value and the limit of detection of the test (the number of cycles performed). The lower the expected Ct value relative to the limit of detection, the greater the probability of detection. For a milk sample from one animal with a Ct value of 16.9 (the population mean peak Ct value), the expected Ct value in a milk sample from 40,000 cattle was estimated to be 33.4 (90% rt-PCR test efficiency), 32.8 (95% rt-PCR test efficiency), and 32.2 (100% efficiency). These values are all well below realistic limits of detection (e.g. 38 cycles [6], 40 cycles [7], 45 cycles [3]). Even at higher Ct values — either due to an infection with lower viral load or a sample time not at the peak — the expected Ct value is likely within realistic limits of detection, particularly for test efficiencies of 90% or more [10]. The expected Ct values would decrease with a greater number of infected cattle making a positive detection even more likely. This suggests that frequent (e.g. weekly) pooled testing of samples from dairy farm milk vats would rapidly identify outbreaks, with a single index case detectable even on large dairy farms (e.g. 30,000–40,000 cattle).

**Figure 3:**
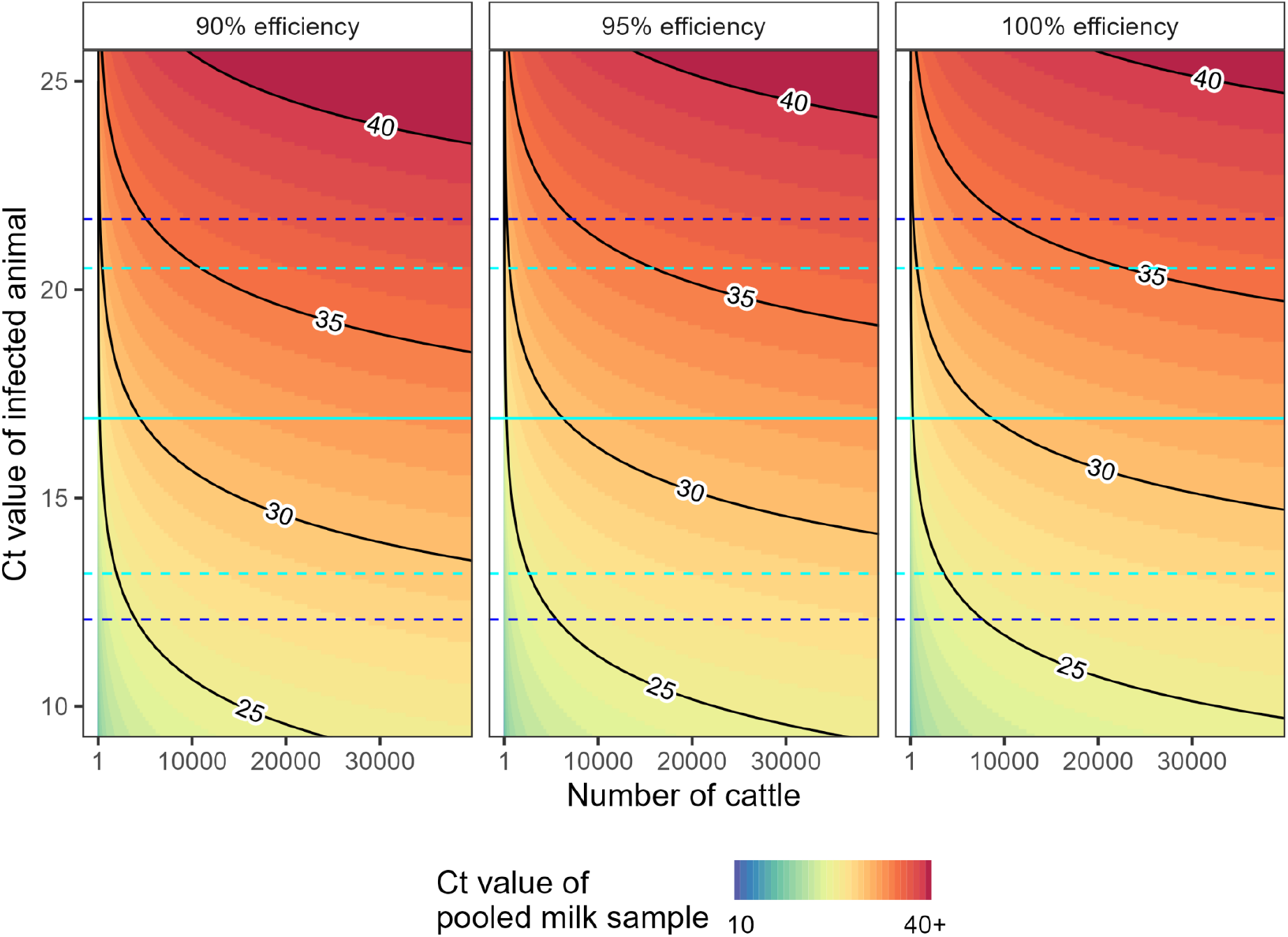
Expected Ct values of pooled milk samples. Expected Ct value of a pooled milk sample (colour) as a function of: the Ct value of the infected animal (y-axis, one infected animal contributing to the pooled sample); the total number of cattle contributing to the sample (x-axis); and the efficiency of the rt-PCR test used (panels). Note that an rt-PCR test of 100% efficiency doubles the viral concentration with each round of amplification (Ct value increasing by 1). Contour lines (Ct values: 25, 30, 35, 40) connect points on the graph with the same value (labelled black lines). The median (solid line) and 95% credible interval (dashed lines) for the population mean minimum Ct value (μ_*P*_) for a single infected animal is highlighted (cyan). The 95% credible interval (dashed lines) for the population distribution ofminimum Ct values using mean population parameters, 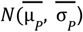, for a single infected animal is also highlighted (blue).

### Relationship between Ct value and infectious virus

We inferred the relationship between Ct value and the log-titre of infectious virus in milk in infected dairy cattle by fitting a statistical model describing a sigmoidal function to a dataset of infected cattle for which both measurements were available. As Ct value decreases — representing a higher viral load — the log-titre of infectious virus increases (Fig.2b). Our estimates suggest that this increase occurs rapidly over a small range of Ct values centered on a critical threshold value of 21.5 (20.1, 23.6), with Ct values above this threshold representing little-to-no infectious viral load. Note that no longitudinal data are available describing Ct value and log-titre of infectious virus for infected cattle over time, and so a hierarchical approach to this part of the analysis could not be taken. Hence our estimates can not account for variation between animals in the relationship between Ct values and log-titre of infectious virus.

### Duration of infectiousness

The dominant route of cow-to-cow transmission is thought to be through infectious virus in an infected animal’s milk leading to direct infection of the mammary gland of another animal, either due to contamination of the environment (e.g. floor and bedding) or contamination of milking apparatus [3,11,12]. The log-titre of infectious virus in milk is thus a useful proxy for estimating the infectiousness of an animal. We have estimated population mean and variation in Ct value trajectories, and the relationship between Ct value and log-titre of infectious virus. We combine these estimates to infer the duration in which infected cattle have a Ct value below the critical threshold value (when they are likely infectious).

Accounting for parameter uncertainty in both models, we estimate that the mean duration of infectiousness of an infected animal is 6.2 (2.8, 13.1) days (Fig.2C). This estimate is sensitive to the critical threshold Ct value, with a higher critical threshold Ct value leading to a longer mean duration of infectiousness (SFig.3). We estimate that there is variation in the duration of infectiousness between animals with median parameter estimates (Fig.2C) suggesting that 3.1% of animals do not become infectious, and that 2.5% of animals are infectious for greater than 17.8 days. We estimate that 95% of cattle will no longer be infectious from 16.6 (8.4, 38.0) days post infection, which has implications for determining the appropriate policy for isolation duration.

### Effectiveness of isolating infected cattle

Using the full posterior for our estimates of the duration of infectiousness (population distribution), we estimated the proportion of transmission events that would be prevented given case isolation (Fig.4, SFig.4). We calculate this reduction in transmission for different proportions of infections that are isolated (0 to 1) and at different times post infection (0 to 15 days). We estimated high uncertainty in the effectiveness of case isolation. For example, if 100% of cases were isolated one week post infection we estimated 32% (6%, 64%) of onward transmission would be prevented. As expected, the longer the estimated duration of infectiousness, the more effective we estimated case isolation to be. This is because longer infectious periods (for the same overall transmissibility and timing of isolation) provide a longer time window for the prevention of onward transmission events.

The effectiveness of case isolation for preventing onward transmission depends on: the proportion of infections that can be detected and isolated; and the relative timing of detection to infectiousness (Fig 4). These two factors are dependent on the rate and timing of clinical symptoms in infected animals, and the ability for symptoms to be recognised (e.g. reduced feed intake and rumination time may be difficult to identify). In some experimentally infected cattle changes to milk color and consistency began as early as two days post inoculation [7] suggesting that careful monitoring of milk output during an outbreak may allow early detection of infections. However, it is currently unclear what proportion of naturally infected cattle exhibit identifiable symptoms and if the timing is comparable to experimentally infected cattle. Epidemiological investigations of the first H5N1 outbreaks on dairy farms reported attack rates of clinically affected cattle ranging from 5–23% [3], but it is unclear what the underlying infection attack rates were for these outbreaks (i.e. including both clinical and subclinical infections). Our results indicate that case isolation is unlikely to be effective if infected animals enter isolation beyond seven days post infection even if a very high proportion (e.g. 90–100%) of infections can be detected and isolated. Further, case isolation within seven days of infection is only likely to be effective if the true duration of infectiousness is within the upper bound of our estimated distribution (e.g. 95% CrI of mean duration of infectiousness is 2.8–13.1 days). In general, higher proportions of and faster rates of infection detection will result in greater effectiveness of case isolation.

**Figure 4:**
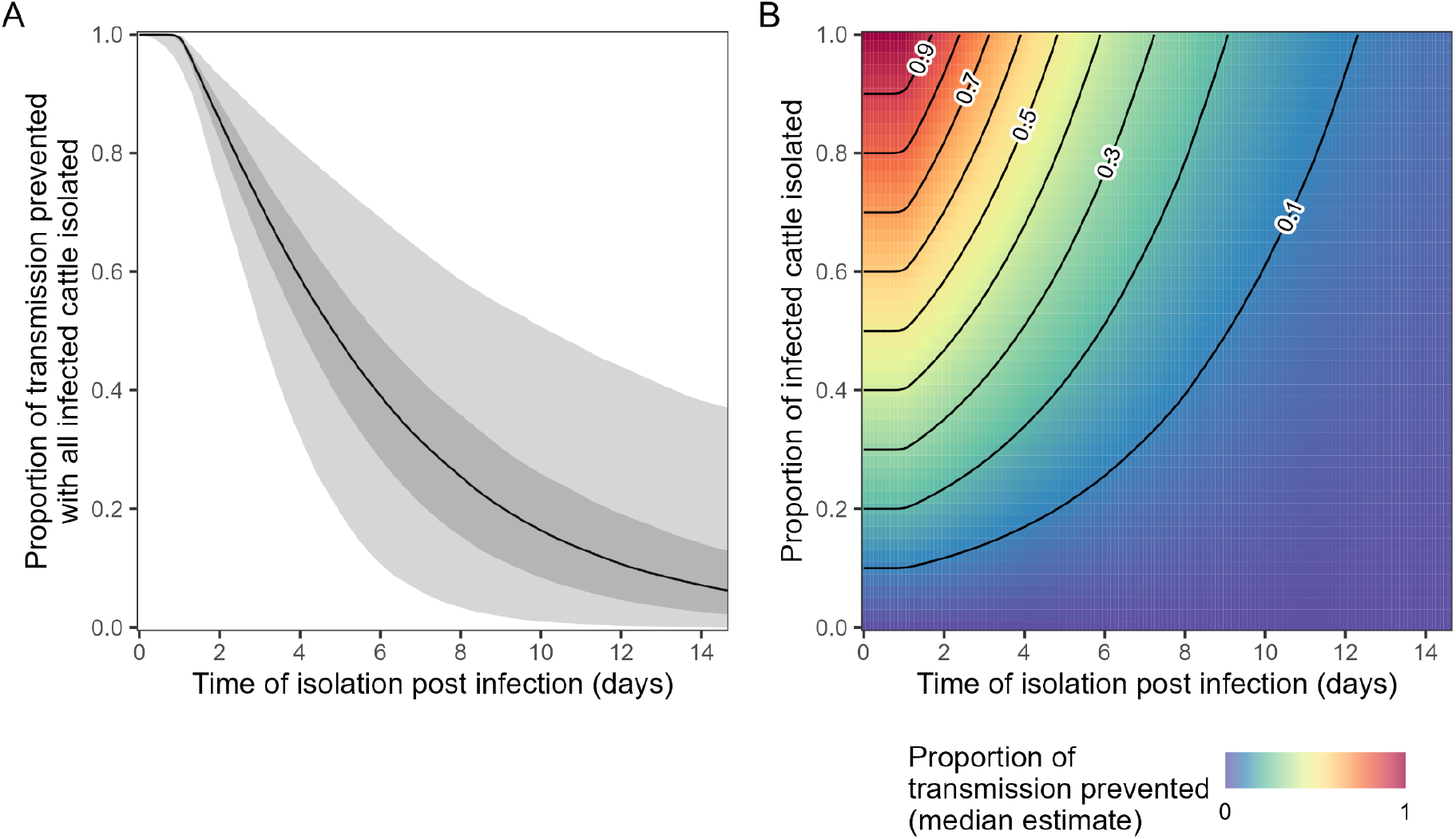
Proportion of transmission prevented by case isolation. **(A)** The proportion of transmission prevented, if all infected cattle are isolated, as a function of the time post infection at which infected cattle are isolated (x-axis). Posterior estimates are shown with median (black line), and 50% (dark shaded region) and 95% (light shaded region) credible intervals. **(B)** The proportion of transmission events prevented (colour) as a function of: the time post infection at which infected cattle are isolated (x-axis); and the proportion of infected cattle that are isolated (y-axis). The proportion of transmission events prevented is calculated across the entire posterior distribution for both models; plotted is only the median value (50 percentile panel). The central 50% and 95% credible intervals for all x- and y-value are shown in SFig.4, and for the case when 100% of infected cattle are isolated, in panel A . Contour lines (proportion of transmission prevented: 0.1–0.9 in intervals of 0.1) connect points with the same value (black lines).

### Limitations

There may be issues with the representativeness of the data, given the small number of animals sampled and the lack of data from naturally infected animals. The only longitudinal data that measured Ct values over the entire course of infection was from experimentally infected cattle; viral kinetics of H5N1 infection in naturally infected cattle may differ. For example, naturally infected cattle may have been initially exposed to a lower volume of infectious virus than experimentally infected cattle (a high volume of infectious virus is often used during experimental inoculation to ensure infection), which could result in a longer time until the peak Ct value is reached [13]. Differences in the time to peak Ct value would alter the period for which cattle remain infectious and thus alter estimates of the effectiveness of control measures such as case isolation. Although we found no evidence for a difference in the viral decay rates between experimentally and naturally infected cattle, the sample size was small. Additionally, the data we had from naturally infected cattle were identified through early outbreak investigations, which may have been preferentially biased towards more symptomatic or severe cases. This underscores the need for more detailed studies aimed at capturing the full course of infection in naturally infected cattle, with unbiased selection of cattle — longitudinal sampling of randomly selected cattle on a dairy farm during an outbreak could provide such data.

Due to the limited data available we could only implement simple models describing the viral kinetics of H5N1 infection in dairy cattle. We assumed a single parameter described the viral decay rate following the peak in Ct value. However, there may be biphasic decay (e.g. as observed in other virological [14,15] and serological [16,17] datasets) with an initially high decay rate followed by a lower long-term decay rate. This may have implications for our estimates of the duration of infectiousness (and effectiveness of case isolation) as a faster decay rate rate could lead to a reduction in the time Ct values remained below the critical threshold value. Additionally, the lower decay rate and the possibility of longer term shedding of virus may have implications for testing strategies — long-term shedding could lead to positive detection for a period of time following the end of an outbreak that would need to be considered when developing interventions. We also lacked longitudinal data describing the relationship between measured Ct values and the log-titre of infectious virus, and so could not fit a hierarchical model that allows for variation in the relationship between individuals in the population. Individual-level variation could influence estimates of the distributions of the duration and timing of infectiousness. Longitudinal studies that measure Ct value trajectories should additionally collect data on the log-titre of infectious virus so that models can be extended to account for individual-level variation.

## Conclusion

We have inferred the viral kinetics of H5N1 infections in dairy cattle. While our statistical models have been designed to analyse currently available data, it will be important to update our estimates as the situation evolves and additional data are collected. Using our estimated Ct value trajectories for the milk of infected dairy cattle, we have demonstrated that pooled testing of milk samples would likely be highly sensitive at detecting the presence of a single infected animal on even the largest dairy farms (e.g. 30,000–40,000 cattle) or central processing facilities. This supports the recent implementation of the United States Department of Agriculture’s National Milk Testing Strategy [18] which includes: testing of samples from milk silos at national dairy processing facilities to determine the presence of H5N1 at the state level; and bulk tank sampling programs in affected states to identify dairy herds affected by H5N1. Additionally, we have estimated the timing and duration of infectiousness, and demonstrated that case isolation is unlikely to be effective for outbreak control if infected animals enter isolation beyond seven days post infection, even if a very high proportion (e.g. 90–100%) of infections can be detected and isolated. Our estimates will be useful inputs to the development of outbreak management guidelines and future modelling analyses required to inform outbreak response strategies.

## Methods

### Ct value data

We retrieved longitudinal data describing the Ct value in milk samples from five experimentally infected (H5N1 B3.13) dairy cattle (inoculated by intramammary route) [6,7], and 12 naturally infected dairy cattle [3]. All animals included in the study were reported to show clinical signs.

### Ct value data: experimentally infected cattle

The longitudinal data for two experimentally infected (H5N1 B3.13) dairy cattle are available from the supplementary material of Baker et al. 2024 [7]. The Ct values were from rt-PCR testing of milk samples in buckets. Ct values were measured daily up to 23 days post inoculation. The limit of detection was a Ct value of 40 (i.e. up to 40 rounds of amplification). The full study protocol is available in the published article [7].

The longitudinal data for three experimentally infected (H5N1 B3.13) dairy cattle was extracted from Halwe et al. 2024 [6]. The Ct values from rt-PCR testing of milk samples were not available in the supplementary materials and so the values were digitally extracted from the original figure (Fig.3 Halwe et al. 2024 [6]) using PlotDigitizer [19]. This extraction process will have introduced a small amount of additional uncertainty into the Ct values. Ct values were measured daily up to 21 days post inoculation. Measurements for one animal were only available up to 3 days post inoculation, and for another up to 13 days post inoculation due to these animals meeting conditions for humane euthanasia. The limit of detection was a Ct value of 38 (i.e. up to 38 rounds of amplification). The full study protocol is available in the published article [6].

### Ct value data: naturally infected cattle

The longitudinal data for the 12 naturally infected dairy cattle are available from the supplementary material of Caserta et al. 2024 [3]. Ct values were measured using rt-PCR testing of milk samples collected 3, 16, and 31 days post clinical diagnosis for a subset of cattle on farm 3 of the study. Data were available on 15 animals, but we excluded three cattle for which there was either only one data point available or for which all measurements were outside the limit of detection. The limit of detection was a Ct value of 45 (i.e. up to 45 rounds of amplification). The full study protocol is available in the published article [3].

### Ct value trajectory model

We fit a Bayesian hierarchical model quantifying the Ct value trajectories in infected dairy cattle. For each experimentally infected dairy cow, *i*, Ct values are modelled as a function of time since infection, *t*_*inf*_ :

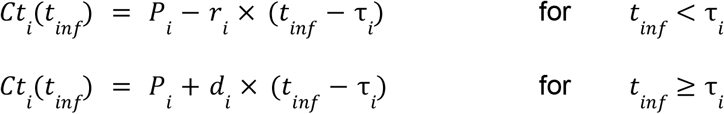

The four parameters are: *P*_*i*_, the minimum Ct value reached; τ_*i*_, the time since infection at which the minimum is reached; *r*_*i*_, the initial viral growth rate (decline in Ct value) before the minimum Ct value is reached; *d*_*i*_, the viral decay rate (increase in Ct value) after the minimum Ct value is reached.

For each naturally infected dairy cow, *j*, Ct values are modelled as a function of time since clinical diagnosis, *t*_*cd*_:

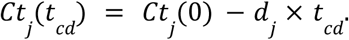

The two parameters are: *Ct*_*j*_(0), the Ct value on the day of clinical diagnosis; and *d*_*j*_, the viral decay rate (increase in Ct value) after the minimum Ct value is reached. This explicitly assumes that the naturally infected dairy cattle were all in the phase of viral decay when their first sample was collected (3 days post clinical diagnosis). This is consistent with our estimates of τ_*i*_ which was 1–2 days post inoculation in experimentally infected cattle. Note that clinical diagnosis will be later than the date of infection as well.

The model was implemented in STAN [20] and parameter posteriors were sampled using a No-U-Turns Sampler [21] fitting to all data (sensitivity analysis described below). The model was run for 4000 iterations across four chains with a burn in period of 1000 iterations. The prior distributions used for all parameters are described in Supplementary Table 1. We assumed that uncensored Ct value data (not at the limit of detection) was normally distributed around modelled Ct values with standard deviation, . Ct value data at the limit of detection were treated as additional parameters with constant priors (outside the limit of detection) and were also assumed to be normally distributed around modelled Ct values with standard deviation, σ_*Ct*_.

The individual-level parameters describing the Ct value on the day of clinical diagnosis for the naturally infected cattle, *Ct*_*j*_(0), were given an uninformative constant prior (with a minimum possible value of 0). All other individual-level parameters were assumed to follow a population-level distribution with parameters describing their mean, and standard deviation, σ:

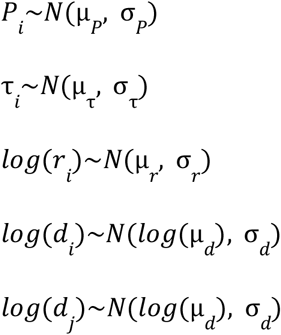

The rate parameters were assumed to be log-normally distributed while the other parameters were assumed to be normally distributed. All individual-level parameters, and population mean and standard deviation parameters, were given uninformative constant priors. Individual-level and population mean viral decay and growth rate parameters were assumed to be in the range 0.01 to 100 per day. Individual-level and population mean minimum Ct value parameters were assumed to be greater than 0. Individual-level and population mean time until minimum Ct value parameters were assumed to be in the range 0 to 10 days post inoculation.

### Sensitivity analyses

We ran additional models as a sensitivity analysis (SFig.1). We fit the model to data from only either experimentally infected cattle or naturally infected cattle (note that for the latter only the viral decay rate parameters can be fit using such data). Finally, we fit the model to all data, but assumed that the viral decay rate parameters for experimentally (‘exp inf’) and naturally (‘nat inf’) infected cattle came from distinct distributions:

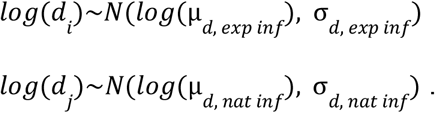

### Pooled testing analysis

We calculated the expected Ct value that would be measured when testing pooled samples of milk in which a single positive milk sample was present. We calculated the expected Ct value of a pooled sample using the equation:

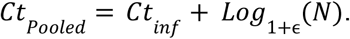

Here *Ct*_*inf*_ is the expected Ct value of the infected animal’s milk sample tested in isolation, ϵ is the PCR efficiency of the rt-PCR test [10], and *N* is the total number of cattle in the sample (*N*-1 are uninfected). This equation assumes that the volume of milk from each cow present in the sample is equal, which may not always be the case as some infected cattle have been reported to have reduced milk output. This equation assumes that when the number of cattle in the pooled sample doubles the viral concentration halves, and so more rounds of amplification are required to detect the virus. Note that when the efficiency of the rt-PCR test is 100% then doubling the number of cattle means an additional round of amplification is required (i.e. the expected Ct value increases by 1).

### Ct value and infectious virus log-titre model

We fit a statistical model to quantify the relationship between Ct value and the log-titre of infectious virus in milk samples. Data on measurements of Ct value and the log-titre of infectious virus from milk samples of infected cattle was available from the supplementary material of Caserta et al. 2024 [4]. Measurements of Ct value (Fig.1a Casera et al. 2024 [4]) and log-titre of infectious virus (Fig.1f Casera et al. 2024 [4]) were matched by sample IDs (see Data and Code availability). In total we were able to identify 43 pairs of measurements. The log-titres of infectious virus were expressed as the *log*_10_ of the 50% tissue culture infectious dose (*TCID*_50_) per millilitre (*TCID*_50_*ml*^−1^). The limit of detection was 10^1.05^*TCID*_50_*ml*^−1^. The full study protocol is available in the published article [3].

We modelled the log-titre (*Log*_10_(*TCID*_50_*ml*^−1^)) of infectious virus, *LT*, as a sigmoidal function of the Ct value:

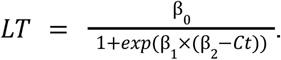

Where β_0_ describes the maximum value of *LT*. β_1_ describes the steepness of *LT*′*s* transition from 0 to its maximum value of β_1_.β_2_ describes the Ct value at *LT* which reaches half of its maximum value. When fitting the model (see below) the posterior distribution for β_1_ only allows for rapid transitions from 0 to the maximum value of LT (i.e. large values of β_1_); the parameter β_2_ thus represents a critical threshold Ct value (referred to as such in the main text) above which the log-titre of infectious virus is zero, and below which the log-titre of infectious virus is at its maximum.

The model was implemented in STAN [20] and parameter posteriors were sampled using a No-U-Turns Sampler [21] fitting to all data (sensitivity analysis described below β_1_). The model was run for 10000 iterations across four chains with a burn in period of 2000 iterations. We allowed for uncertainty in both measured Ct values and log-titres of infectious virus by assuming modelled *LT* was a a function of modelled Ct values, and treating modelled Ct values (each corresponding to a data point) as additional parameters with uninformative constant prior distributions (limited to be positive in value). For the eight points in which the log-titre of infectious virus was at the limit of detection, we treated the true value of the log-titre as an additional parameter with a constant prior distribution and upper limit of 1.05 *Log*_10_*TCID*_50_*ml*^−1^ (the limit of detection). We assumed that Ct value data were normally distributed around the corresponding modelled Ct value with standard deviation, σ_*Ct*_. We assumed that log-titre data were normally distributed around the corresponding modelled *LT* value with standard deviation, σ_*LI*_. The parameters σ_*Ct*_, and σ_*LI*_ were assumed to have uninformative constant prior distributions. Parameters, β_0_, β_1_ and β_2_ were similarly given uninformative constant priors: β_0_ was limited to be strictly positive; β_1_ was limited to therange -100 to 0; and β_2_ was limited to the range 0 to 30.

### Estimating the duration of infectiousness

Posterior model estimates of the relationship between Ct value and log-titre of infectious virus found a critical threshold Ct value (β_2_) at which the log-titre of infectious virus rapidly increased from 0 to a maximum. Under the assumption that a high log-titre of infectious virus in milk samples corresponds to an animal being infectious, we estimate the duration and timing of infectiousness by estimating the duration in which individuals in a population have a Ct value below the critical threshold value.

For each set of population parameters (means and standard deviations) from the posterior distribution of the Ct value trajectory model, we randomly draw a set of individual level parameters and simulate an individual’s Ct value trajectory (1000 trajectories simulated for each set of parameters). For all individuals we calculate the duration the Ct value remains below the critical threshold value for each value of β_2_ in the posterior distribution (log-titre of infectious virus to Ct value model). This gives us the distribution for the duration of infectiousness across individuals for each set of population parameters. We calculate the mean duration of infectiousness for each set of parameters, which gives us our posterior distribution for the mean duration of infectiousness. Similarly, we can calculate the time at which 95% of individuals are no longer infectious for each set of parameters. We report the median and 95% credible intervals of each distribution in the main text.

We also estimate the posterior distribution of the mean duration of infectiousness, assuming that β_2_ is known exactly to highlight how our estimate depends on the parameter value (SFig.3). We also estimate the distribution for the duration of infectiousness using the mean values of the population parameters from the posterior distribution of the Ct value trajectory model (Fig.2C).

### Effectiveness of case isolation

As before, for each set of population parameters (mean and standard deviations) from the posterior distribution of the Ct value trajectory model, we simulate individual Ct value trajectories (1000 trajectories simulated for each set of parameters). We then calculate the proportion of the population infectious (Ct value below threshold value, accounting for entire posterior distribution of β_2_), as a function of time since infection, for each set of parameters.

We then calculate the proportion of infections prevented for different values of: the time post infection at which infected cattle are isolated (0–15 days post infection considered); and the proportion of infected cattle that are isolated (0–1 considered). We calculate the proportion of infections prevented by each pair of case isolation values for each set of population parameters giving us the posterior distribution for the proportion of infections prevented for each pair of case isolation values. We then plot the median, and central 50% and 95% credible intervals of the posterior distribution for all case isolation values considered (Fig.4).

## Data and code availability

The data and code required to reproduce these analyses is available at https://github.com/Eales96/H5N1_viral_kinetics (Zenodo DOI: https://doi.org/10.5281/zenodo.1478844)

## Acknowledgements

OE is supported by a University of Melbourne McKenzie Fellowship. FMS is supported by the National Health and Medical Research Council of Australia through the Investigator Grant Scheme (Emerging Leader Fellowship, 2021/GNT2010051). JMM is supported by the Australian Research Council through the Laureate Fellowship Scheme (FL240100126). FMS and JMM’s research is also supported by an Australian Research Council Discovery Project Grant (DP240102286). This research is supported by the Australian Consortium of Epidemic Forecasting and Analytics (ACEFA), a National Health and Medical Research Council of Australia Centre of Research Excellence (2035303).

## Supplementary figures

**Supplementary figure 1:**
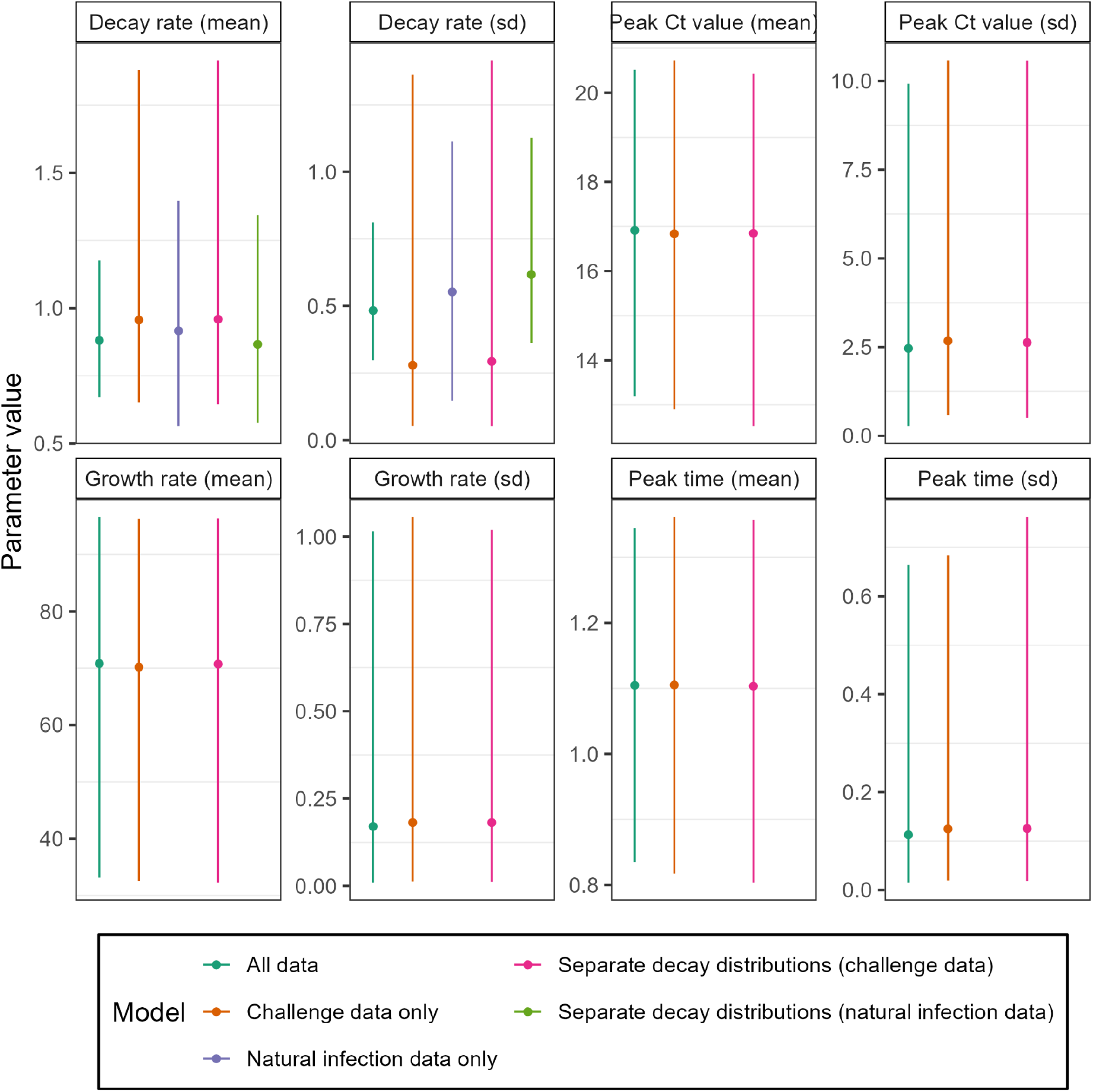
Sensitivity of Ct value trajectory model parameter estimates. Median (point) and 95% credible intervals (lines) for posterior parameter estimates for the Ct value trajectory model using different subsets of data. Models include: fitting to all data, the main model used for analysis (dark green); fitting only to the data for experimentally infected cattle (orange); fitting only to the data for naturally infected cattle (purple); and fitting to all data, but assuming separate distributions for the viral decay rate between experimentally and naturally infected cattle (pink, green for decay distribution of naturally infected cattle). Note that when fitting the model only to naturally infected cattle only the viral decay rate distribution can be estimated.

**Supplementary figure 2:**
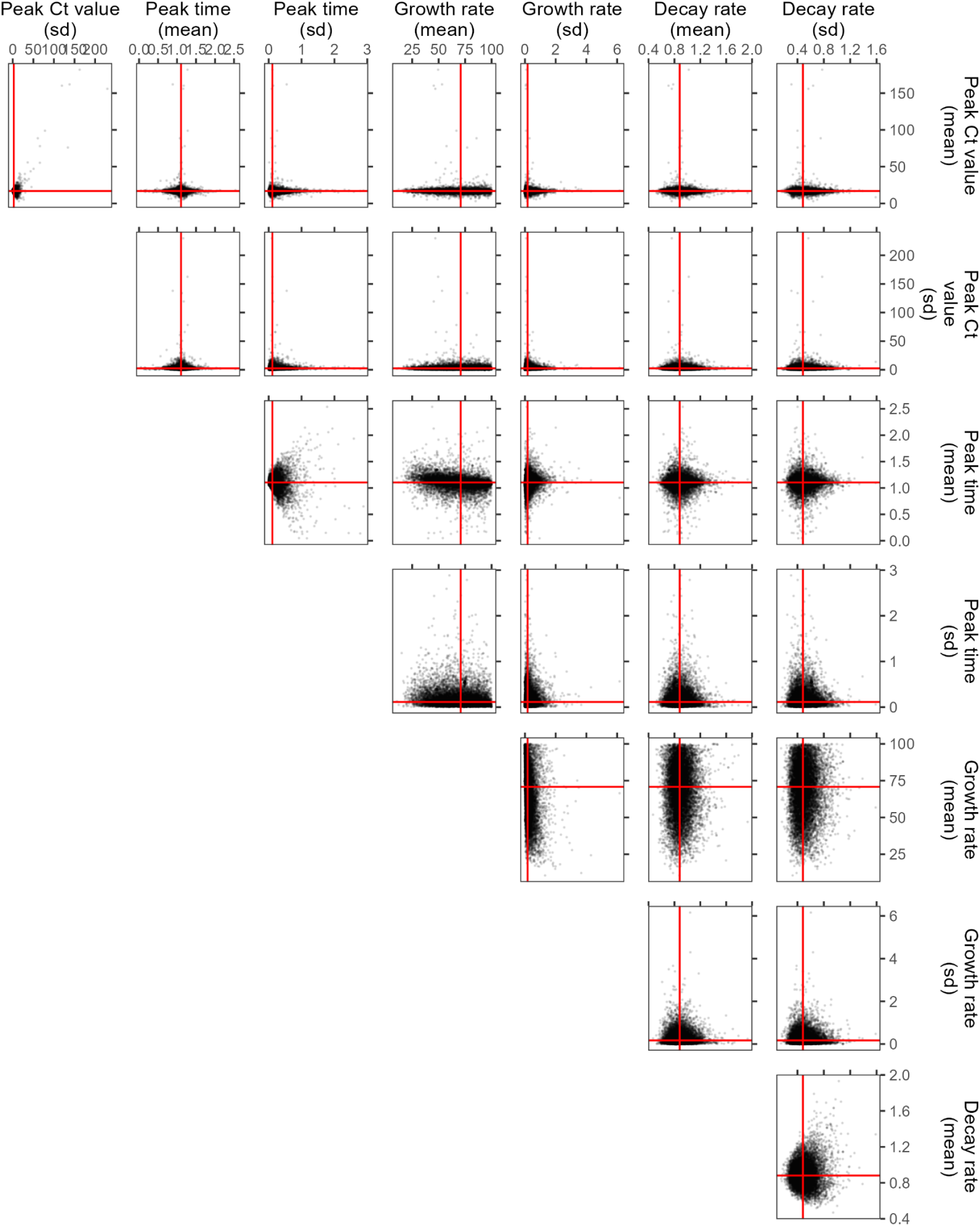
Posterior distributions of Ct value model population parameters. Posterior samples (black points) from the Ct value model for all population parameters (means and standard deviation). The distribution of posterior samples is shown for each pair of population parameters. Also shown is the median of the posterior distribution for each individual parameter (red lines). These median parameter values are also shown in SFig.1 and were used in Fig.2a to plot the 95% interval of the estimated population distribution of Ct value trajectories (using median parameter estimates).

**Supplementary figure 3:**
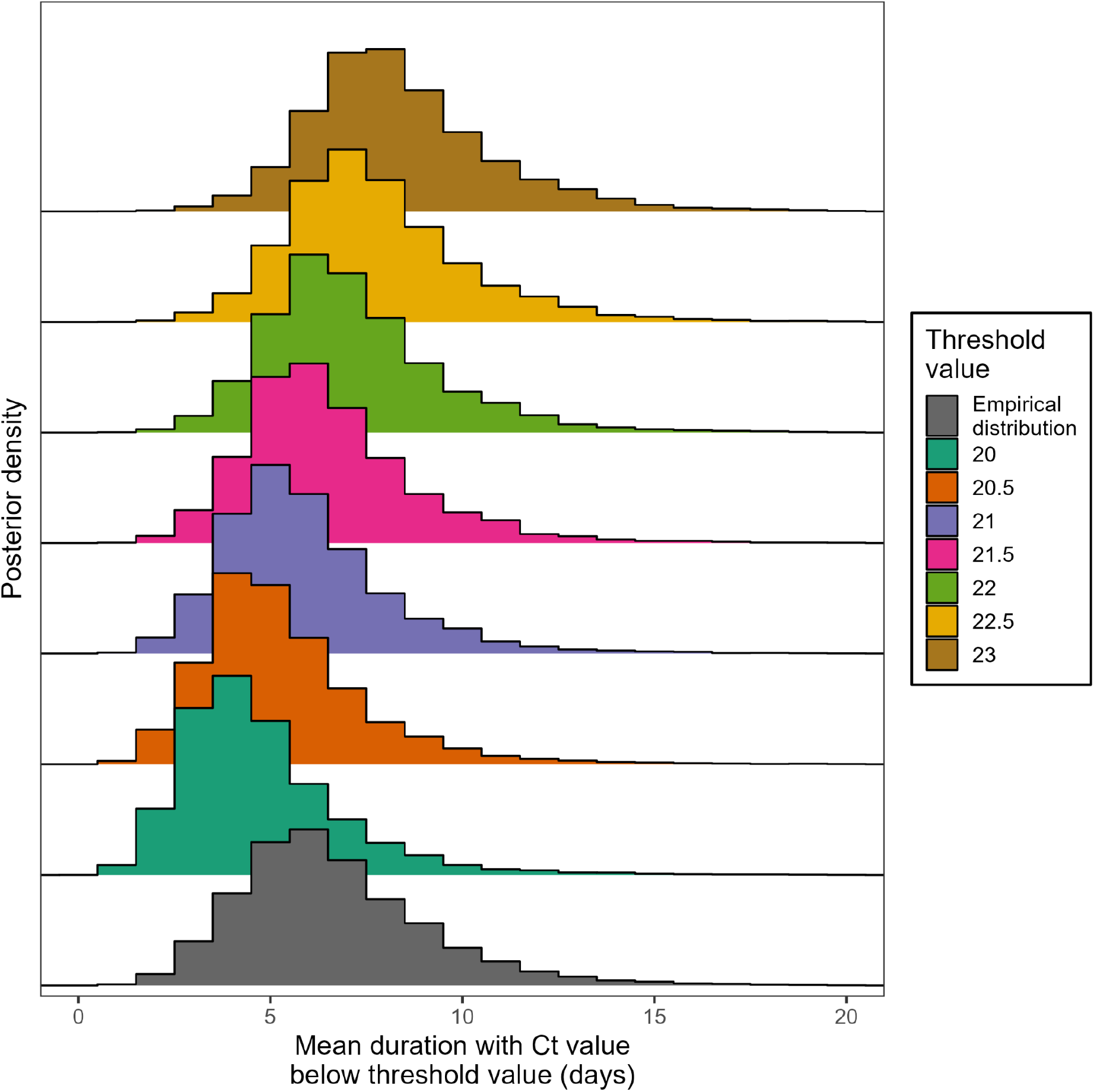
Posterior distribution for the mean duration of infectiousness. The posterior distributions for estimates of the mean duration for which cattle’s Ct value remains below a threshold value. Shown are the posterior distributions for different choices of the threshold value (colours). Also shown is the posterior distribution using the empirical distribution of the threshold value (as estimated using the model linking Ct value and the log-titre of infectious virus, parameter). Ct values above the estimated threshold value represent little-to-no infectious virus and so are unlikely to be infectious. This posterior distribution thus represents our posterior distribution for the duration of infectiousness.

**Supplementary figure 4:**
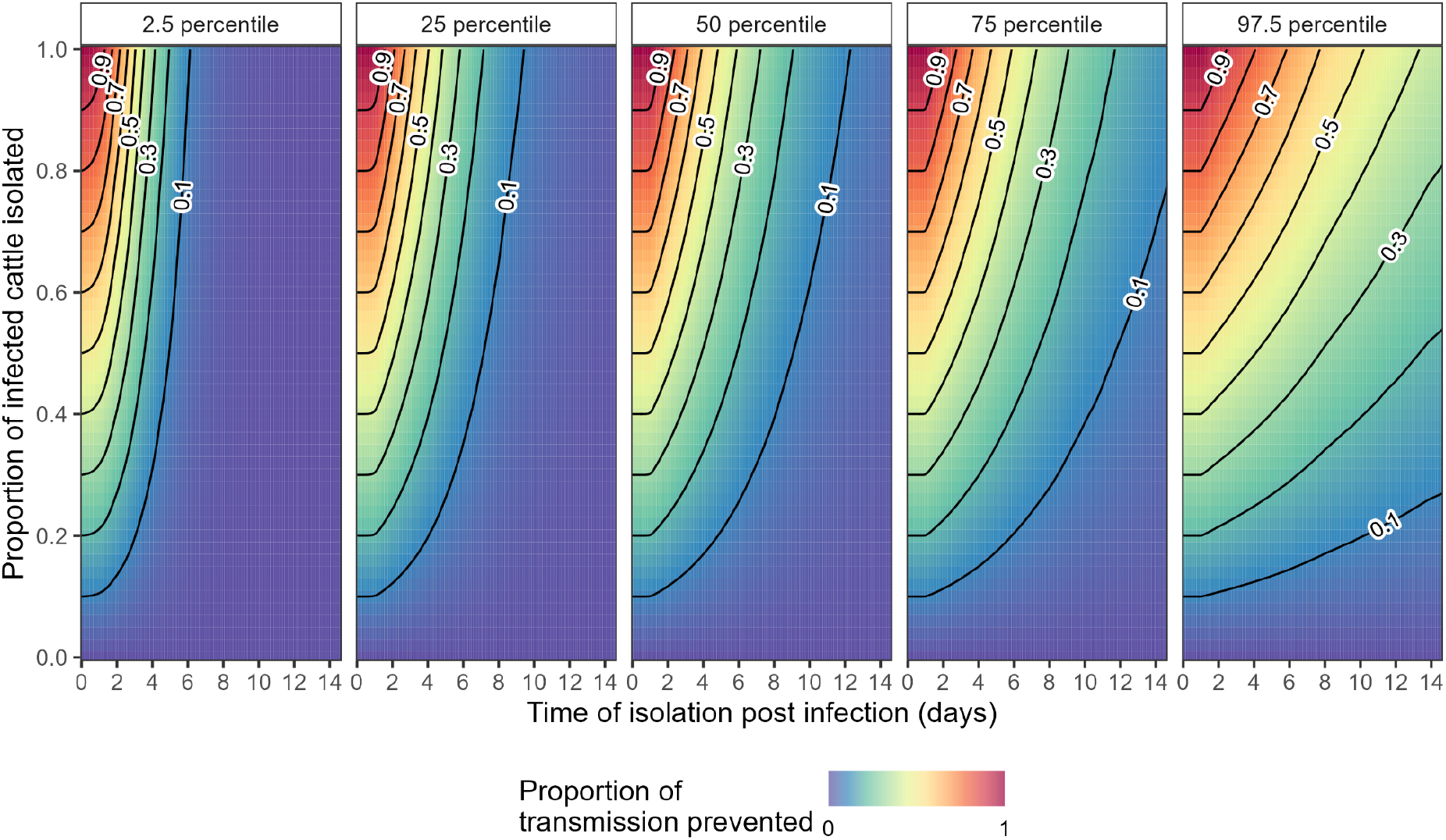
Proportion of infections prevented by case isolation. The proportion of transmission events prevented (colour) as a function of: the time post infection at which infected cattle are isolated (x-axis); and the proportion of infected cattle that are isolated (y-axis). The proportion of transmission events prevented is calculated across the entire posterior distribution for both models; plotted is the median value (50 percentile panel), and the central 50% (25–75 percentile panels) and 95% (2.5–97.5 percentile panels) credible intervals. Contour lines (proportion of transmission prevented: 0.1–0.9 in intervals of 0.1) connect points on each panel with the same value (black lines).

