## Supplementary Table 1 for "Viral kinetics of H5N1 infections in dairy cattle"

| Parameters | | Prior distribution | Minimum | Maximum |
|---|---|---|---|---|
| **Individual-level parameters** | | | | |
| **Experimentally infected dairy cattle (i)** | | | | |
| P_i | Minimum Ct value reached | P_i ~ Normal(mu_P, sigma_P) | 0 | - |
| tau_i | time since infection at which minimum is reached | tau_i ~ Normal(mu_tau, sigma_tau) | 0 | 10 |
| r_i | viral growth rate (before minimum Ct is reached) | log(r_i) ~ Normal(log(mu_r), sigma_r) | 0.01 | 100 |
| d_i | viral decay rate (after minimum Ct is reached) | log(d_i) ~ Normal(log(mu_d), sigma_d) | 0.01 | 100 |
| **Naturally infected dairy cattle (j)** | | | | |
| Ct_j(0) | Ct value on day of clincial diagnosis | Uninformative constant prior | 0 | - |
| d_j | viral decay rate | log(d_j) ~ Normal(log(mu_d), sigma_d) | 0.01 | 100 |
| **Population-level parameters** | | | | |
| mu_P | Mean of distribution for P_i | Uninformative constant prior | 0 | - |
| sigma_P | Standard deviation of distribution for P_i | Uninformative constant prior | 0 | - |
| mu_tau | Mean of distribution for tau_i | Uninformative constant prior | 0 | 10 |
| sigma_tau | Standard deviation of distribution for tau_i | Uninformative constant prior | 0 | - |
| mu_r | Mean of distribution for r_i | Uninformative constant prior | 0.01 | 100 |
| sigma_r | Standard deviation of distribution for r_i | Uninformative constant prior | 0 | - |
| mu_d | Mean of distribution for d_i and d_j | Uninformative constant prior | 0.01 | 100 |
| sigma_d | Standard deviation of distribution for d_i and d_j | Uninformative constant prior | 0 | - |
| **Data parameters** | | | | |
| sigma_Ct | Standard deviation of Ct value data around modelled values | Uninformative constant prior | 0 | - |
| Censored_Ct | Censored Ct value data were treated as an addtional parameter | Constant prior | Limit of detection | - |
